# Rice rhizospheric microbes confer limited Arsenic protection under high Arsenic conditions

**DOI:** 10.1101/2020.11.02.365312

**Authors:** Victoria Gundlah-Mooney, Harsh P. Bais

**Affiliations:** Department of Plant and Soil Sciences; Delaware Biotechnology Institute, University of Delaware, Newark, Delaware; AP Biopharma, 590 Avenue 1743, Newark, DE 19713

**Keywords:** Rice, Rhizospheric microbes, microbiome, Arsenic

## Abstract

Rice (*Oryza sativa*) is a staple food crop worldwide and plays a critical role in ensuring food security as the global population continues to expand exponentially. Groundwater contamination with Arsenite [As(III)], a naturally occurring inorganic form of arsenic (As), leads to uptake and accumulation within rice plants. As a result, grain yield is lowered, the overall plant health is diminished, and there is a risk of arsenic toxicity from grain consumption. It was previously shown that a novel bacterial strain from the rice rhizosphere may reduce As accumulation in rice plants exposed to low levels of environmental As. We hypothesized that different rice varieties may exhibit varying responses to high As levels, resulting in differences in As uptake and toxicity. Utilizing the natural rice rhizospheric microbes, we initiated a set of hydroponic experiments with two rice varieties, Nipponbare (As tolerant) and IR66 (As susceptible). Rice varieties exposed to high As(III) concentration (50 μM) showed changes in both aboveground and belowground traits. As-tolerant Nipponbare varieties show grain production at high As(III) concentrations compared to the As-susceptible IR66 variety. Supplementation of natural rice rhizospheric microbes as single inoculums showed varied responses in both As-tolerant and As-susceptible varieties. Three natural rice rhizospheric microbes *Pantoea* sps (EA106), *Pseudomonas corrugata* (EA104), and *Arthrobacter oxydans* (EA201) were selected based on previously reported high Iron (Fe)-siderophore activity and were used for the hydroponic experiments as well as a non-rice rhizospheric strain, *Bacillus subtilis* UD1022. Interestingly, treatment with two strains (EA104 and EA201) led to reduction in As(III) uptake in shoots, roots, and grains and the degree of reduction of As(III) was pronounced in As-susceptible IR66 varieties. Non-rice rhizospheric UD1022 showed subtle protection against high As toxicity. High As(III) treatment led to lack or delay of flowering and seed setting in the As-susceptible IR66 variety. The data presented here may further the understanding of how beneficial microbes in the rhizosphere may help rice plants cope with high concentrations of As in the soil or groundwater.

## Introduction

Millions of people depend on rice to meet their daily nutritional requirements. It is consumed by more people than any other agricultural crop (Bouman, et al., 2015; Yadav and Kumar, 2019). A rice shortage would not only increase pricing but could also lead to malnutrition and starvation for those who depend on it as a staple food source. Increasing rice production and yields will not only help meet the demand for food, but also regulate the cost. Developing innovative solutions to maximize yields and reduce crop loss is an active field of research (Kumarathilaka et al., 2018; Ansari et al., 2015).

There are many biotic and abiotic stressors that impact rice production and yields. Abiotic stressors affecting production are climate changes, drought, fertilizers, and heavy metal contamination, with As being a particular threat to human health (Sytar et al., 2019). Arsenic (As) is a naturally occurring metalloid found throughout the world, contaminating both soil and water sources (Sharma et al., 2014). Plants grown in contaminated soil or irrigated with contaminated water can suffer from heavy metal toxicity. Heavy metals can be taken up by plants and stored in the roots, stems, leaves, fruit, and grains which are subsequently consumed by animals or people (Raessler et al., 2018). Constant consumption of contaminated food and water can lead to serious health problems such as, dermal lesions, gastrointestinal and cardiovascular issues, neurological deterioration, or cancer (Das and Sarkar, 2017).

In addition to impacting human health, As toxicity also affects plant growth and yield. Depending on the toxic element, plants can display symptoms such as, but not limited to, inhibited seed germination, chlorosis, decreased photosynthetic efficiency, browning of leaves and roots, inhibited growth, decreased water uptake, decreased nutrient uptake, reduced yields, and plant death (Moulick et al., 2018). One of the crops most affected by As toxicity is rice due to several factors. Geographically, As contamination is common in the same locations where rice is predominantly grown (Kumarathilaka et al., 2018). Additionally, the semi-aquatic growth requirements of rice lead to both soil and water contamination throughout the rice growth period (Sharma et al., 2014).The concentration of As in the rice grains, which are the primary source of human consumption and ingestion, can differ among rice cultivars. Some rice cultivars such as Nipponbare have been found to be more tolerant to As contamination while others are more susceptible (ref). The most toxic and prevalent forms of As are the inorganic arsenate [As(V)] and arsenite [As(III)] (Awasthi et al., 2017). Redox potential is a big factor in which form of As is found in the ground water and surrounding sediments (Kumarathilaka et al., 2018). As(V) is abundant in surface waters that are more oxygenated. The reduced form, As(III), is more abundant in groundwater and can be sixty times more toxic and generally more available in paddy soil than As(V) (Sharma et al., 2014 and Seyfferth et al., 2017).

Transport of As from paddy soil to rice grains is not fully known. However, it is shown that inorganic species are less mobile in the plant than organic As species (Awasthi et al., 2017; Hashimoto and Kanke, 2018). In fact, it is predicted that 10% of As(III) will reach the shoot and 3.3% of As(III) will reach the grains (Zhao et al., 2012). It is well accepted that inorganic As species enter rice roots due to structural similarities they share with the phosphorous and silicic acid transport pathways. The structure of As(V) is analogous to inorganic phosphate. Whereas As(III) mimics the structure of silicic acid and is thought to use the low silicon 1 (Lsi1) and low silicon 2 (Lsi2) silica transporter pathways to enter the plant (Seyfferth et al., 2017). The Lsi1 protein is located on the outer side of the plasma membrane. It is a passive Aquaporin channel permeable to both silica and As. The Lsi2 protein is an active transporter polarly located to the inner side of the plasma membrane (Sun et al., 2018). Both are highly expressed in the exo and endodermis of rice root cells where casparian strips are formed and nutrient uptake is regulated (Seyfferth et al., 2017; Sun et al., 2018). As (III) is the dominant form of As loaded into the xylem. Their strategic locations are vital for As transport to the xylem and into the rest of the plant (Sun et al., 2018). It is not fully understood how other biotic components in soil, including microbes, facilitate or abate As uptake in plants.

The rhizosphere is the defined region of soil surrounding a plant’s roots, which is influenced by root exudates and the soil microbiome (Osman et al., 2017). The microbiome consists of consortium of diverse microbial species found in the soil rhizosphere (Lakshmanan, et al., 2015). Previous studies have shown that on a whole, the microbiome functions to positively influence the plants they surround (Osman et al., 2017). The beneficial bacteria found in the rhizosphere are termed plant growth promoting rhizobacteria (PGPR) (Lakshmanan et al., 2015; Osman et al., 2017). They are known to promote nutrient acquisition, positively influence growth and development, influence plant physiology and metabolism, protect against pathogens, and play a role in immune response and tolerating abiotic stressors (Osman et al., 2017; Busby et al., 2017). For example, previous studies where bacteria were isolated from the rhizosphere of M-104 rice, a japonica cultivar grown abundantly in California, have shown a beneficial response in rice against both biotic and abiotic stress (Lakshmanan et al., 2015; Spence et al., 2014). The experiments speculated that microbes found and associated with field rice may offer more protection than bioinoculants from another plant species due to their ability to survive and compete with other agents in the rice rhizosphere (Spence et al., 2014). In this study, both natural rice rhizospheric microbes and a gram-positive, soil associated *Bacillus subtilis* UD1022 strain were used to evaluate if bioinoculants mitigate the impacts and accumulation of As toxicity in rice. These strains were selected based on their ability to increase iron (Fe)-siderophore activity (Lakshmanan et al., 2015). It is known that both Fe and As compete in the soil (Gustave et al., 2018). Thus, using microbes that mobilize Fe in soil may abate As uptake. Strains such as *Pantoea* sp. EA106 has been shown to increase iron siderophore in culture (Lakshmanan et al., 2015). This is important because siderophore production positively influences Fe uptake, therefore benefiting plant growth (Lakshmanan et al., 2015). Strains such as EA104 a *Pseudomonas corrugata* strain., have been found to produce antimicrobial secondary metabolites and are used as biocontrol bacteria (Spence et al., 2014). The EA201 strain, an *Arthrobacter* sp., is a gram-positive bacterium commonly found in soil. Previous studies have shown *Arthrobacter* may be useful for bioremediation of contaminated soils (Dsouza et al., 2015). The non-rice rhizospheric strain UD1022, a *Bacillus subtilis* strain, has shown to be a potent PGPR involved in elevating plant response against multiple stress regimes (Zheng et al., 2018). Previous studies have shown that *B. subtilis* can produce surfactants inducing antifungal activity (Lakshmanan et al., 2015). Looking at each isolate individually in association with As can help us further understand their potential to abate As uptake and As toxicity in rice plants.

The primary goal of this work is to identify phenotypical changes in As-resistant and susceptible rice varieties exposed to high As(III) concentrations. The phenotypic changes were recorded for both aboveground and belowground portions of the plant, post exposure to a high concentration of As(III). This was done using a tritrophic model encompassing [As(III)], rice rhizospheric microbes, and the plant as a platform for testing phenotypic outcomes and functional responses in plants exposed to both biotic and abiotic components. Having shown that few rice rhizospheric microbes may abate As toxicity under low As environment, we tested plant response supplemented with rice microbiome exposed to high As environment. We also hypothesized that the plant response in As-tolerant and susceptible varieties may vary +/− microbe/As supplementation. We employed phenotypic and molecular assays to evaluate the bacteria – bacteria interactions *in vitro* and *in vivo* on the plant root. Together, these results may provide the basis for future studies to refine the hypothesis that the beneficial microbiome can be used to ameliorate As uptake in rice plants.

## Materials and Methods

### Set up of Hydroponic rice plants

#### Preparing Rice Seeds

The outer husk layer of each rice seed was removed, and each seed was inspected for maturity and fungus. Seeds that appeared mature and fungus-free were added to a 50 mL conical tube, in a biosafety cabinet, and soaked in 30 mL of 50% bleach solution for ten minutes with occasional swirling. After ten minutes the bleach was drained into a beaker and the seeds were washed with autoclaved deionized water three times. Standard petri dishes were prepared with one piece of autoclaved chromatography paper and 3.5 mL of autoclaved deionized water. Sterilized seeds were transferred to the prepared petri dishes using sterile forceps. Five to seven seeds were added to each dish before sealing the plates with parafilm and labeling the dish with seed type, name, and date. Labeled petri dishes were placed under growth lights in the lab and set to a 16-hour photoperiod for seven to ten days until germinated. This procedure was used to prepare all rice seeds used in the hydroponic experiments for both As-tolerant Nipponbare and As-susceptible IR66 rice varieties.

#### Hydroponic conditions for rice growth

After the rice seeds had germinated, the seedlings were transferred to the hydroponic nursery system in the greenhouse growth chambers. Before transfer, foam plugs (Identi-Plug® Plastic Foam Plugs, Jaece) were prepared by cutting them in half once, cutting a slit into one side, and then autoclaving. Once all the supplies were prepared, germinated seeds and supplies were transferred to the campus greenhouse. Seedlings were individually removed from the petri dishes and inserted into a foam plug. The plug, with seedling, was then inserted into an opening in the hydroponic nursery system filled with rice nutrient solution from the International Rice Research Institute (Yadav and Kumar, 2019). The rice seedlings were in the nursery system for 7 days before being transferred to the large hydroponic buckets.

After seven days, the hydroponic buckets were prepared to transfer the rice seedlings. To prepare the 2-gallon bucket, a 1^1/4^-inch hole was first drilled into the center of the lid. The hydroponic buckets were filled with one liter of 8x concentrated rice nutrient media (Yadav and Kumar, 2019) and filled the rest of the way with water. Buckets were then transferred into the growth chamber with a growth cycle of 14 hr light (28°C, 70% RH), and 10 hr dark (26°C, 60% RH). A rice seedling was taken from the nursery system and carefully inserted into the hole of the lid and placed on the 2-gallon bucket. For each experiment, eight replicates per treatment group were used.

### Supplementation of bacteria and arsenic to hydroponic rice cultures

#### Preparing bacterium for inoculation

Under sterile conditions a loop of bacteria was removed from the working glycerol stock of the bacteria of interest *Pantoea sps* (EA106), *Pseudomonas corrugate* (EA104), *Arthrobacter oxydans* (EA201), *Bacillus subtilis* (UD1022). Bacteria was cultured in LB in a 30°C incubator (EA106/UD1022 16-24 h, EA104/EA201 24-48 h). Once the bacteria had grown, in a sterile hood, a single colony was selected and transferred to a flask with LB liquid media. The flask was put into a 30°C shaking (200 rpm) incubator overnight.

After the bacteria culture was grown, 45 mL of culture was distributed into 50 mL conical tubes and centrifuged down (4°C, 3000 rpm) to a pellet with an initial spin of 25 minutes. The supernatant was poured off and the pellet was washed with deionized water and then spun down for 10 minutes (4°C, 3000 rpm). The pellet was washed two additional times. Next, 10-15 mL of deionized water was added to each conical tube and the pellet was resuspended. The resuspended liquid culture from each tube was added to an autoclaved flask and the volume was brought to 200 mL using more deionized water. The cell count and OD were determined using a spectrophotometer (BIORAD Spectrophotometer). The number from the spectrophotometer was used to calculate the amount of culture needed to inoculate 1 × 10^6^ cell/mL into the appropriate hydroponic cultures.

#### Preparing Arsenic for experimental treatment

A stock solution of 0.01 M sodium (meta) Arsenite (NaAsO_2_) was made to be used in all lab experiments containing As for a final As (III) concentration of 5 μM NaAsO_2_. As was added 48 hours post bacterial treatment in rice hydroponic cultures. For high toxicity experiments, plants were irrigated with the rice nutrient solution containing 5 μM As, leading to final concentration of As reaching to 50 μM.

#### Plant materials, growth materials and conditions

*Oryza sativa* Nipponbare seeds were taken from stocks bulked in Bais lab at University of Delaware. *Oryza sativa* IR66-103-2 seeds were obtained from the National Small Grains Collection in Aberdeen Idaho and then bulked in the greenhouse. All plants were grown in the growth chambers in Fischer Greenhouse with a daily cycle of 14 hr light (28°C, 70% RH), and 10 hr dark (26°C, 60% RH).

#### Imaging and harvest

Hydroponic plants were imaged at different stages of growth. Photos were taken of both shoots and roots against a black background, using a representative plant from each treatment group. Once the hydroponic plants reached maturity, plants were harvested for further elemental analysis. The timeline to maturity differed between Nipponbare and IR66 varieties. As the experiments progressed, a plant imaging box was made to take pictures of submerged roots at different stages of development.

#### Microscopy

Root samples from rice plants (IR66) treated with EA106 (1 × 10^6^ cells per mL) with and without As (50 μM) were harvested at days 20 and 21 (96 and 120 hours post microbial inoculation) and stained with SYTO13. Samples were imaged using a Zeiss LSM 710 confocal microscope with a 40 × 1.2 numerical aperture (NA) water immersion objective. Images were analyzed using the Ziess Zen software.

#### As analysis

Quantification of Total As Content Plants grown in an 8-L hydroponic system for ~125 days and treated with As(III), EA104, EA201, UD1022 and EA106 as mentioned above were used for quantification of arsenic. At harvest, the root and shoot samples were collected and dried at 65°C for 3 days and weighed. The total concentration of As in the leaves and seeds was measured by ICP-OES at UD soil testing lab, University of Delaware.

#### Data analysis

Data was presented as mean with standard error. The statistical software JMP11 was used to analyze the data. The data was analyzed using one-way analysis of variance (ANOVA), and post hoc mean separations were performed by Tukey’s HSD test, and results were statistically different when p < 0.05.

## Results

### Supplementation of Pantoea sps. (EA106) in rice hydroponics under elevated As environment

Both the As-tolerant (Nipponbare) and As-susceptible (IR66) rice varieties were treated with *Pantoea* sp (EA106) [hereafter EA106] under high toxic (HT) As[III] (50 μM) environment. In plant treatment groups containing both bacteria and As, plants were pre-treated with EA106 before As exposure for efficient establishment of the microbial inoculum. Plants were treated with EA106 (~10^6^ cells/ml) and As was added in the hydroponic set up post 48h of microbial inoculum treatment. Plants were imaged for aboveground and belowground traits post treatments. Plants were photographed on multiple time points, but the most distinct phenotype in between treatments were observed during 7-8 weeks of growth post-treatment. Plants treated with a high concentration of As showed signs of toxicity and stunted growth at both root and shoot levels (Figure 1). Plants treated with EA106 showed increased growth response compared to As-treated plants (Figure 1). Although the co-inoculation of EA106 and As resulted in no statistical difference in shoot biomass or grain yield (Figure 2, P = 0.95), there was a slight increase in observable shoot growth in between As and EA106 treated plants (Figure 1). Arsenic exposure to IR66 resulted in drastic decrease in biomass and yield. As a matter of fact, none of the IR66 plants exposed to As produced seeds (Figure 1). Nipponbare plants treated with high As displayed signs of abiotic stress in the form of chlorosis and reduced growth (Figure 1). To maintain the highest level of photosynthetic ability, the leaves should be tall and upright, at harvest, the As treated plants showed disorganized shoot structure with droopy leaves (Figure 1). The Nipponbare roots treated with highly toxic levels of As showed reduced biomass (Figure 2). However, root length does not appear to be affected post As treatment (Figure 1). The IR66 rice treated with EA106 bacteria under highly toxic As conditions displayed similar results as Nipponbare plants, though plant shoot organization is not defined until later in developmental phase (Figure 1, harvest: 125 days). The biomass of IR66 roots treated with As showed less dry weight biomass accumulation than the plants treated with As and EA106 bacteria (Figure 1, harvest: 125 days). Nipponbare plants exposed to As + EA106 showed slight decrease in total As content in shoots and roots but not in the grains (Figure 1). This is consistent with Zhoa et al., (2012) paper where the roots contain ~89% of total arsenic, shoots retain ~10%, and grains retained ~3%. The Nipponbare plants had far less As uptake into the tissues than the IR66 plants (Figure 2). This is consistent with the suggestion that the Nipponbare cultivar is tolerant to As. Statistical analyses comparing treatment groups within Nipponbare and IR66 shows no significance in total plant biomass between control and EA106 treatments compared to the As treatments in both rice varieties (Figure 2).

**Figure 1:**
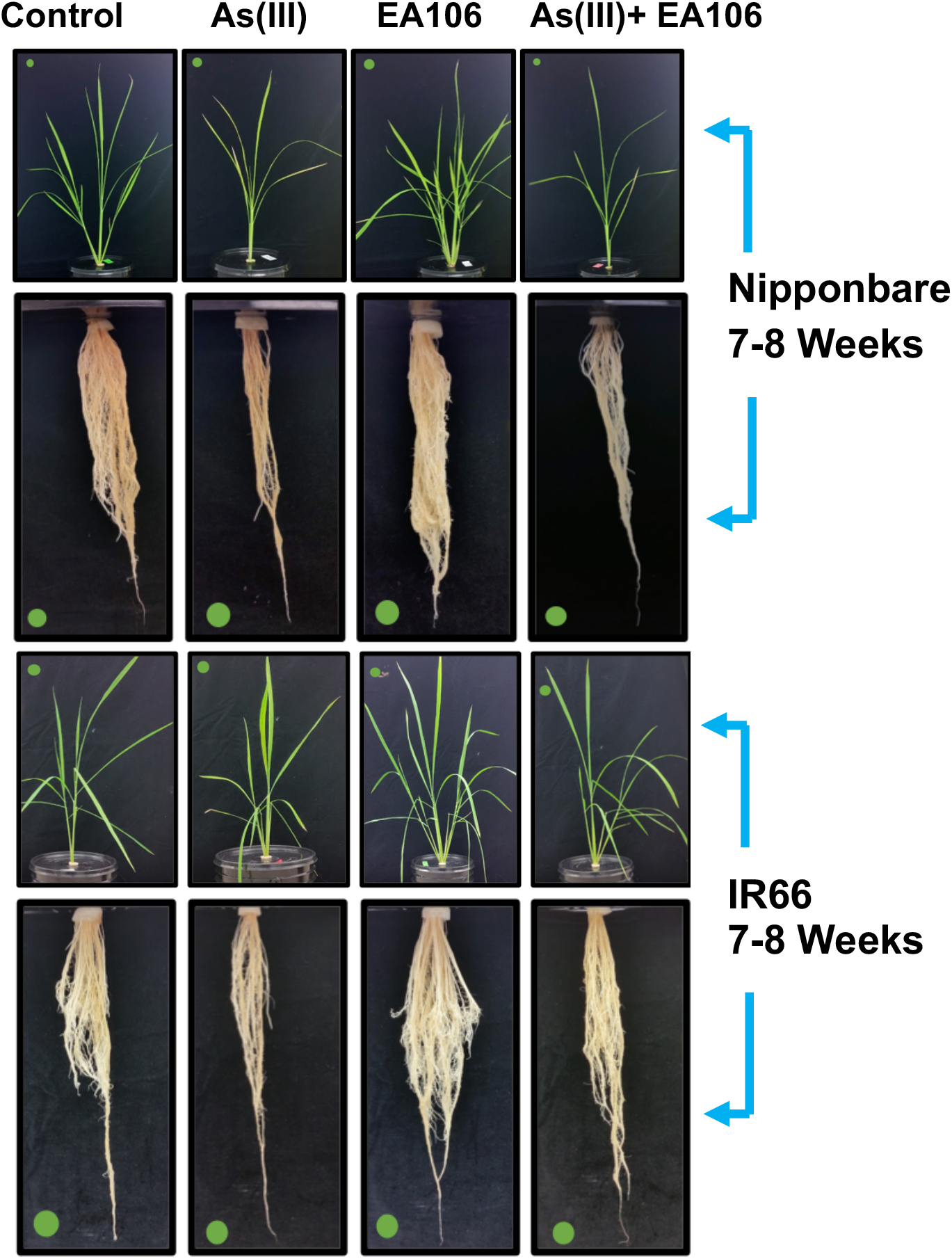
Nipponbare and IR66 rice plants inoculated with *Pantoea* sps. strain EA106 and treated with a high concentration of As[III] (~50 μM). Images show phenotypic differences in the shoots and roots of each plant post +/− EA106/As treatments. Representatives are shown at 7-8 weeks post inoculation and/or arsenic treatment. The green dot scales to 0.75 inches in diameter.

**Figure 2:**
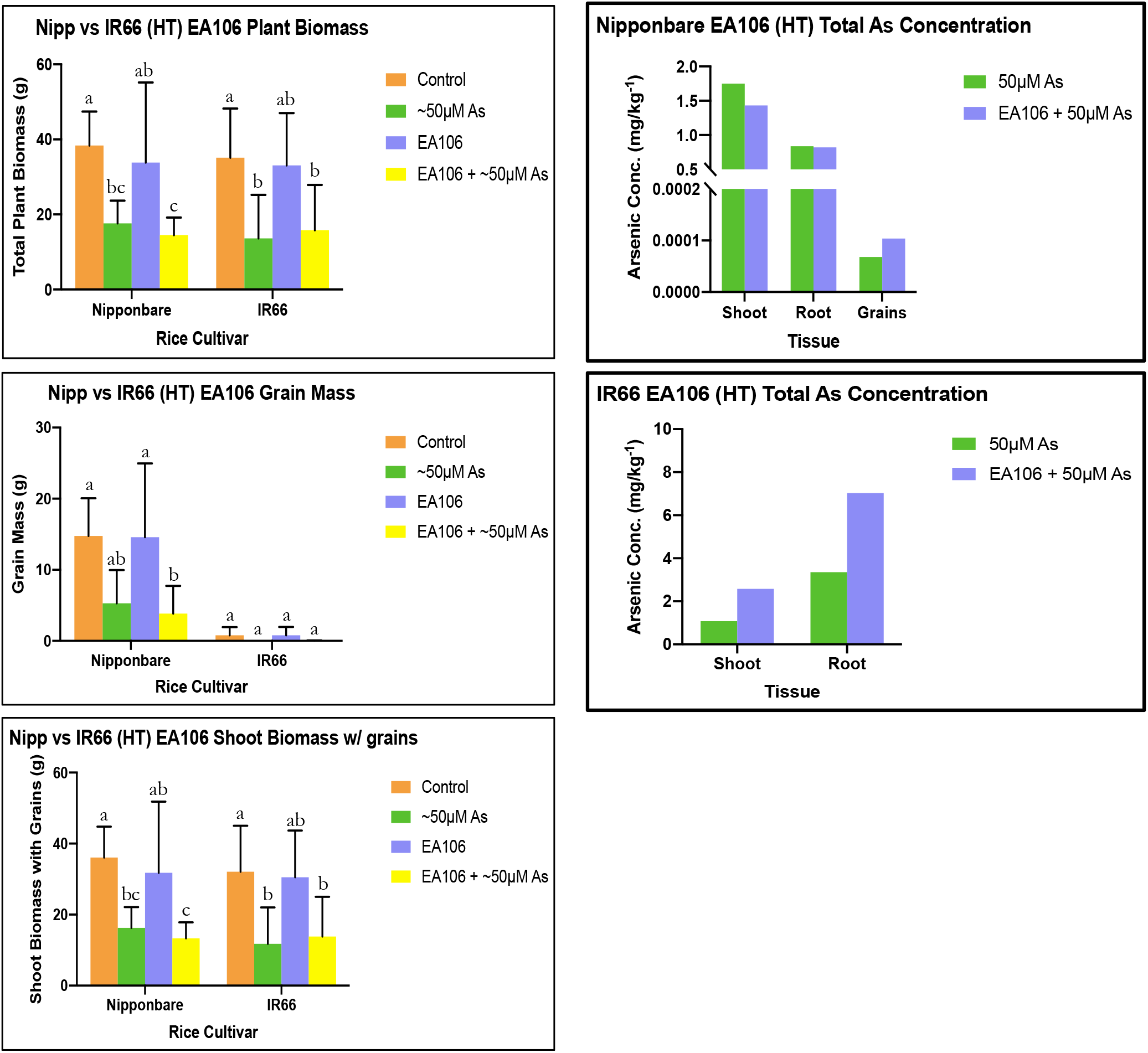
Total plant biomass for Nipponbare and IR66 rice exposed to As [III](~50 μM) and EA106 bacteria. Means of common letters are not significantly different at *P* ≤ 0.05; according to Duncan’s Multiple Range Test.

These results indicate that the addition of the rice rhizospheric microbe EA106 under a high As environment did not increase plant health, though the level of protection that EA106 provided for Nipponbare was more pronounced than IR66. We posited that the lack of protection by EA106 may be due to acute phytotoxicity caused by increased As concentration. The micrographs depicting EA106 colonization +/− As, show reduced EA106 colonization in both root tip and central elongation zone under highly toxic As treatments (Figure S1), suggesting that increased As concentration may also be microbicidal.

### Supplementation of *Pseudomonas corrugate* (EA104) in rice hydroponics under elevated As environment

In contrast to EA106 treatments, rice plants grown in hydroponics treated with *Pseudomonas corrugata* (EA104) showed a plant growth promoting phenotype and better protection against As compared to lone As and control treatments (Figures 3–4). EA104 treated plants displayed more efficient shoot organization from the other treatment groups as well as more established root systems and higher grain production (Figures 3–4). The Nipponbare roots treated with highly toxic levels of As showed a reduced biomass but again no truncation in root length was observed (Figures 3–4). In IR66 rice cultivar treated with EA104 bacteria under high toxicity As conditions, As stress at week seven and eight were visible where it was not as obvious in Nipponbare rice until a later growth stage (Figures 3–4, data not shown). The As treated plants were not as full as the control and EA104 treated plants. In addition, the leaves of As-treated plants showed more chlorosis compared to control and EA104-treated plants and roots looked smaller (Figures 3–4). The biomass of IR66 roots treated with As only, appeared to have less biomass than the plants treated with As and EA104 (Figures 3–4).

**Figure 3:**
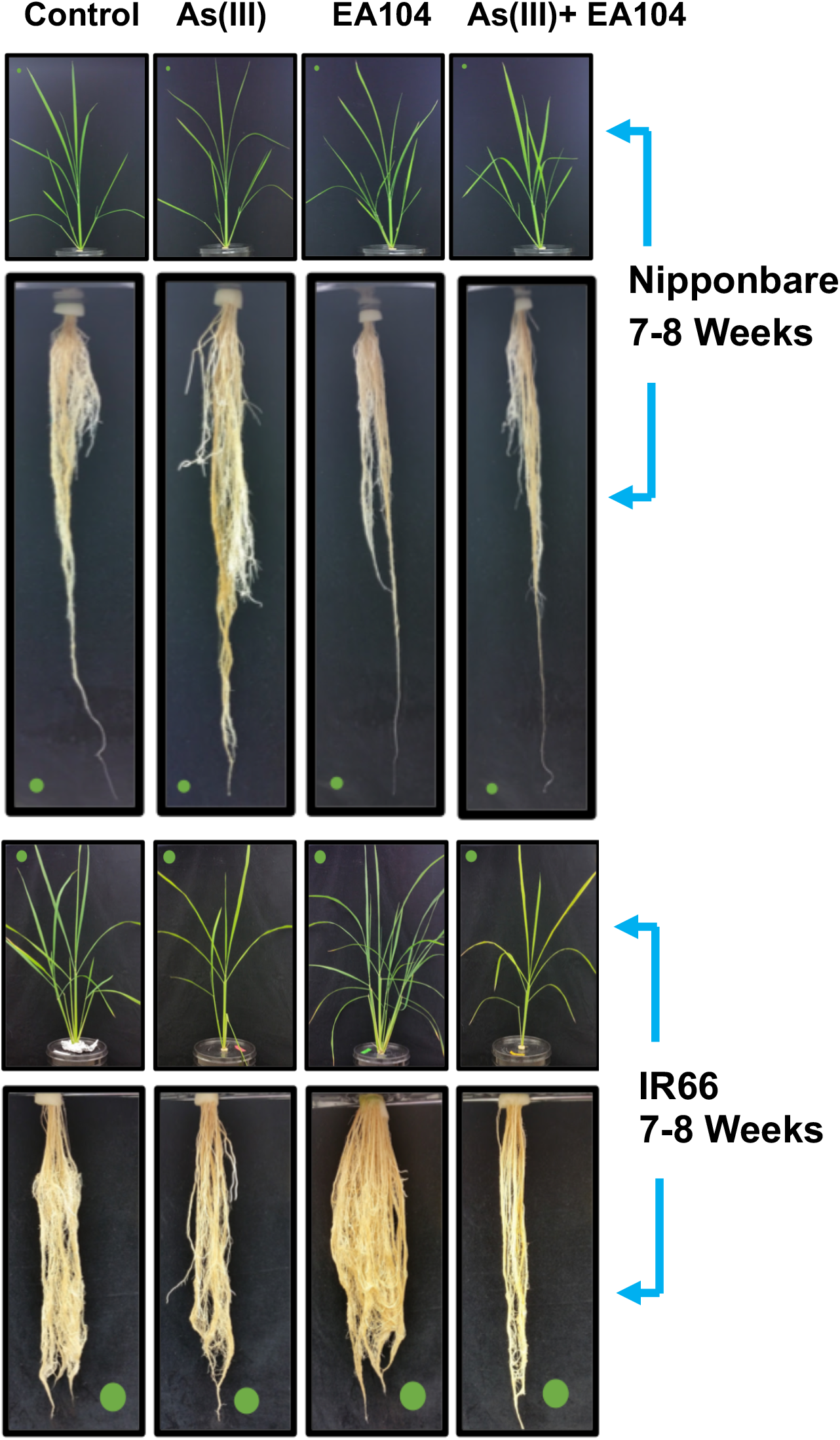
Nipponbare and IR66 rice plants inoculated with EA104 and treated with a high concentration of As (~50 μM). Images show phenotypic differences in the shoots and roots of each plant. Representatives are shown at 7-8 weeks post inoculation and/or arsenic treatment. The green dot scales to 0.75 inches in diameter.

**Figure 4:**
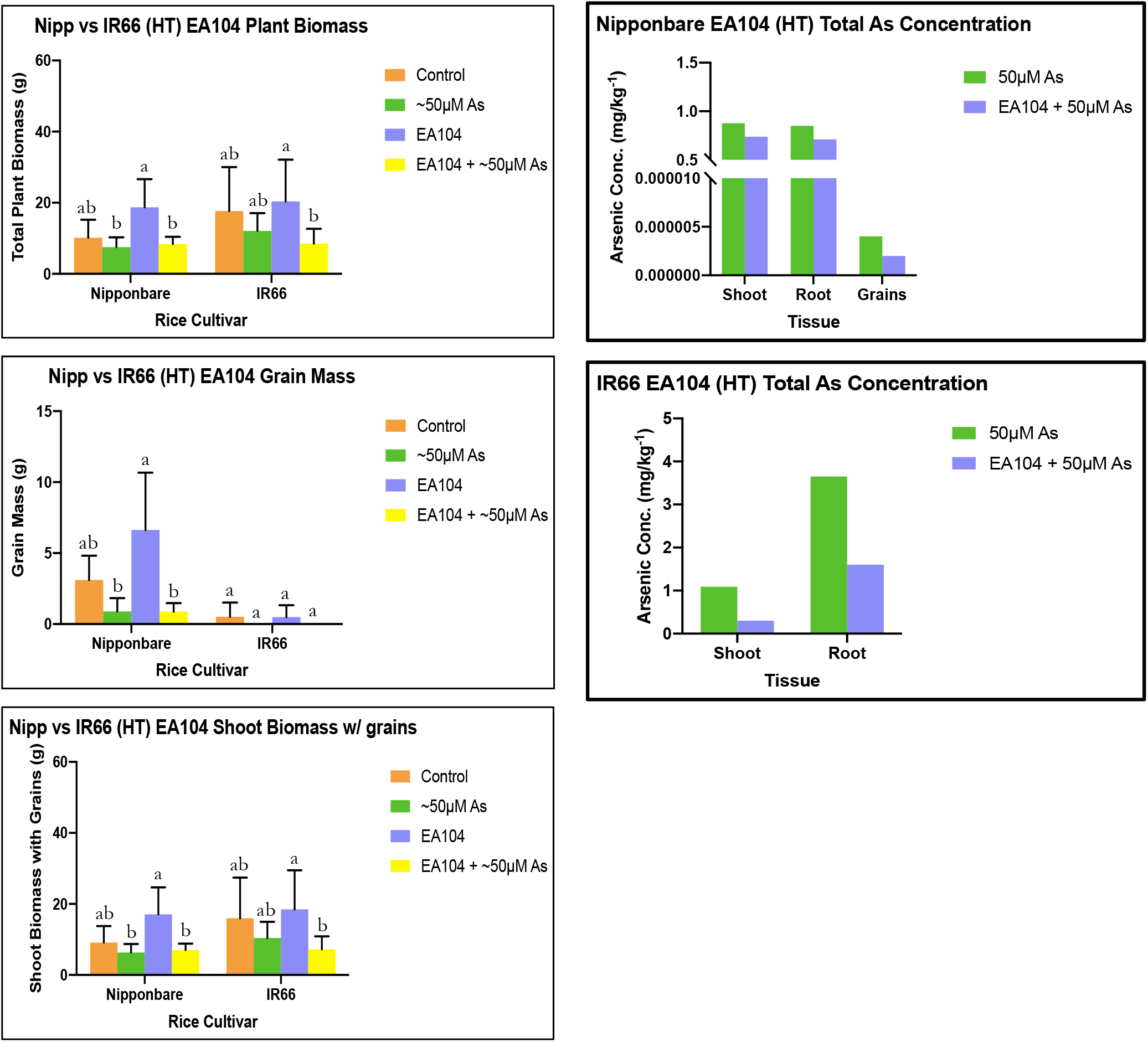
Total plant biomass for Nipponbare and IR66 rice exposed to As [III](~50 μM) and EA104 bacteria. Means of common letters are not significantly different at *P* ≤ 0.05; according to Duncan’s Multiple Range Test.

Statistical analysis for Nipponbare shows significance in total plant biomass for EA104 only bacterial treatment when compared to the other treatment groups (Figures 3–4; P=0.09). In IR66 rice there appears to be significance between the EA104 only bacteria treated group compared to the other treatments (Figures 3–4). Grain mass and yield was higher for all treatment groups in the Nipponbare cultivar with EA104 only being slightly more significant than control and the As treated plants (Figures 3–4). IR66 again had little to no grain yield for all treatments and there was no statistical significance in between the treatment groups (Figures 3–4). Harvest index for Nipponbare EA104 only and control groups was statistically significant over the As treated groups (data not shown). Total shoot biomass shows the EA104 treatment group to be statistically more significant in both rice cultivars (Figures 3–4). Arsenic concentrations in both cultivars yielded results comparable to those for the EA106 high toxicity As treatments. Again, the Nipponbare plants displayed overall lower concentrations of As in the plant tissues than the IR66 cultivar (Figures 3–4). These results are consistent with those from the EA106 high toxicity As treatments further supporting Nipponbare to be an As tolerant cultivar. Both rice varieties treated with EA104 and As showed reduced As in shoots, roots and grains (only for Nipponbare) compared to lone As treatments. The results suggest that EA104 supplementation to rice cultures may provide better protection against high As environment compared to EA106.

### Supplementation of *Arthrobacter oxydans* (EA201) in rice hydroponics under elevated As environment

All the bacterial strains utilized in this study were selected based on the increased Fe-siderophore activity (Lakshmanan et al., 2015). *Anthrobacter oxydans* (EA201), like other two strains (EA106 and EA104) also revealed increased Fe-siderophore activity (Lakshmanan et al., 2015), which may attenuate As uptake in plants under low As (5 μM) treatments. Consistent with other bacteria treatments (EA106, EA104), the untreated control and EA201 treated plants showed an increased biomass compared to the As treated plants (Figures 5–6). In the IR66 rice trial with EA201 bacteria, the phenotype of the As only plant is consistent with other microbial treatments. Interestingly, and unlike other As + microbial inoculation treatments, EA201 plus As treated plants produced grains in IR66 variety (Figures 5–6).

**Figure 5:**
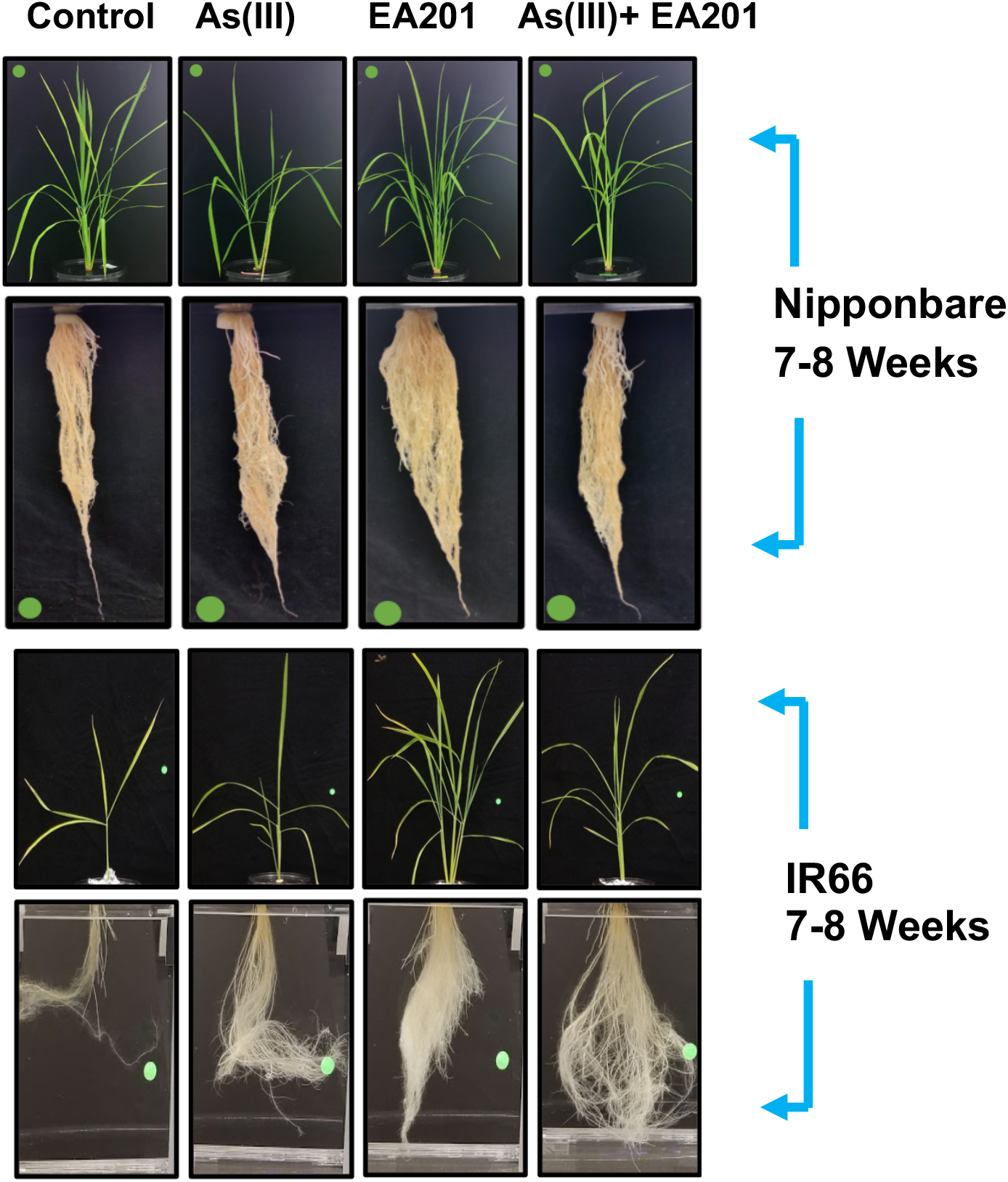
Nipponbare and IR66 rice plants inoculated with EA201 and treated with a high concentration of As (~50 μM). Images show phenotypic differences in the shoots and roots of each plant. Representatives are shown from four treatment groups at 7-8 weeks post inoculation and/or arsenic treatment. The green dot scales to 0.75 inches in diameter.

**Figure 6:**
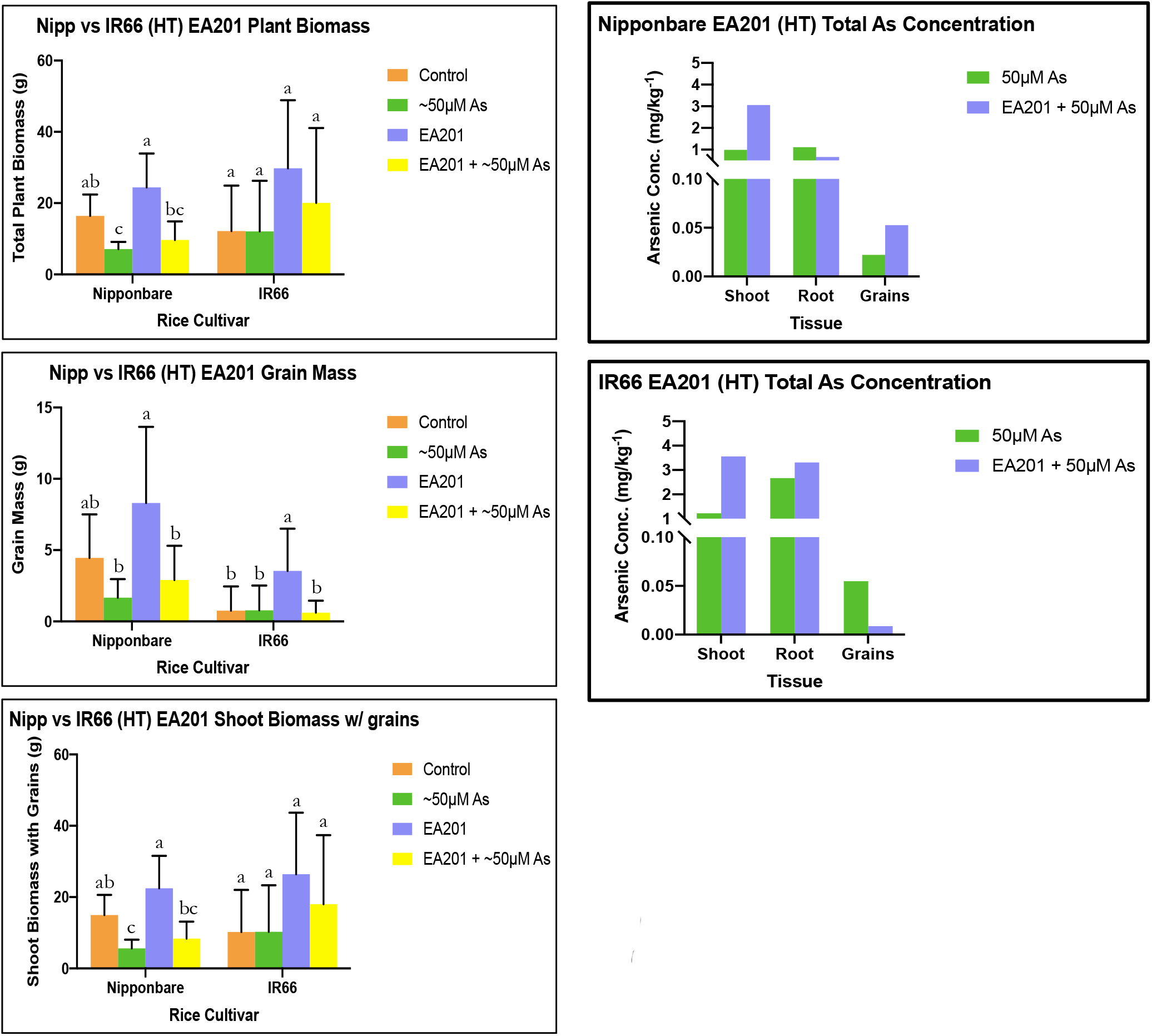
Total plant biomass for Nipponbare and IR66 rice exposed to As [III](~50 μM) and EA201 bacteria. Means of common letters are not significantly different at *P* ≤ 0.05; according to Duncan’s Multiple Range Test.

In both rice varieties, the EA201-only treatment produced significantly more grains than the other treatment groups (Figures 5–6). The harvest index is statistically the same across all treatments for the Nipponbare rice plants but the EA201 only treatment group is significant over the rest for the IR66 variety (data not shown). The progression of As concentrations in the Nipponbare tissues (shoots/roots/grains) is comparable to other Nipponbare and microbial treatments but overall concentration was slightly elevated (Figures 5–6). Interestingly, the total As content in grains in IR66 and Nipponbare varied with the EA201+ As treatments. In the IR66 grain, As content decreased post As treatment compared to Nipponbare plants, suggesting that EA201 may reduce As uptake in the grains of As susceptible rice variety under high As toxic environment.

### Supplementation of *Bacillus subtilis* (UD1022) in rice hydroponics under elevated As environment

All the other bacteria strains (EA106, EA201 and EA104) used in this study originated from the rice rhizosphere and showed increased Fe-siderophore activity (Spence et al., 2014; Lakshmanan et al., 2015). Of the natural rice rhizospheric isolates, EA104 showed the most promising protection against high As concentrations. We also used a non-natural rice rhizospheric isolate, *Bacillus subtilis* (UD1022); UD1022 is used as a classical plant growth promoting rhizobacteria (PGPR) and has shown to impart plant protection against both biotic and abiotic stresses (Bais et al., 2011; Lakshmanan et al., 2015). In the UD1022 treatments, at harvest, all Nipponbare rice plants except those treated with only As produced grains (Figures 7–8). Plants treated with As displayed similar phenotypes as seen in previous bacterial treatments. The root biomass (at 4 weeks) of the control and UD1022 only Nipponbare rice plants was more abundant than the As treated plants (Figures 7–8). IR66 rice plants treated with only UD1022 had a better shoot structure than any other treatment groups. The As only treated plants looked severely inhibited in comparison to UD1022 + As or UD1022 treatments (Figures 7–8). This trend is carried on into the IR66 plant roots, where the As only plants have much less biomass than the other treatment groups (Figures 7–8).

**Figure 7:**
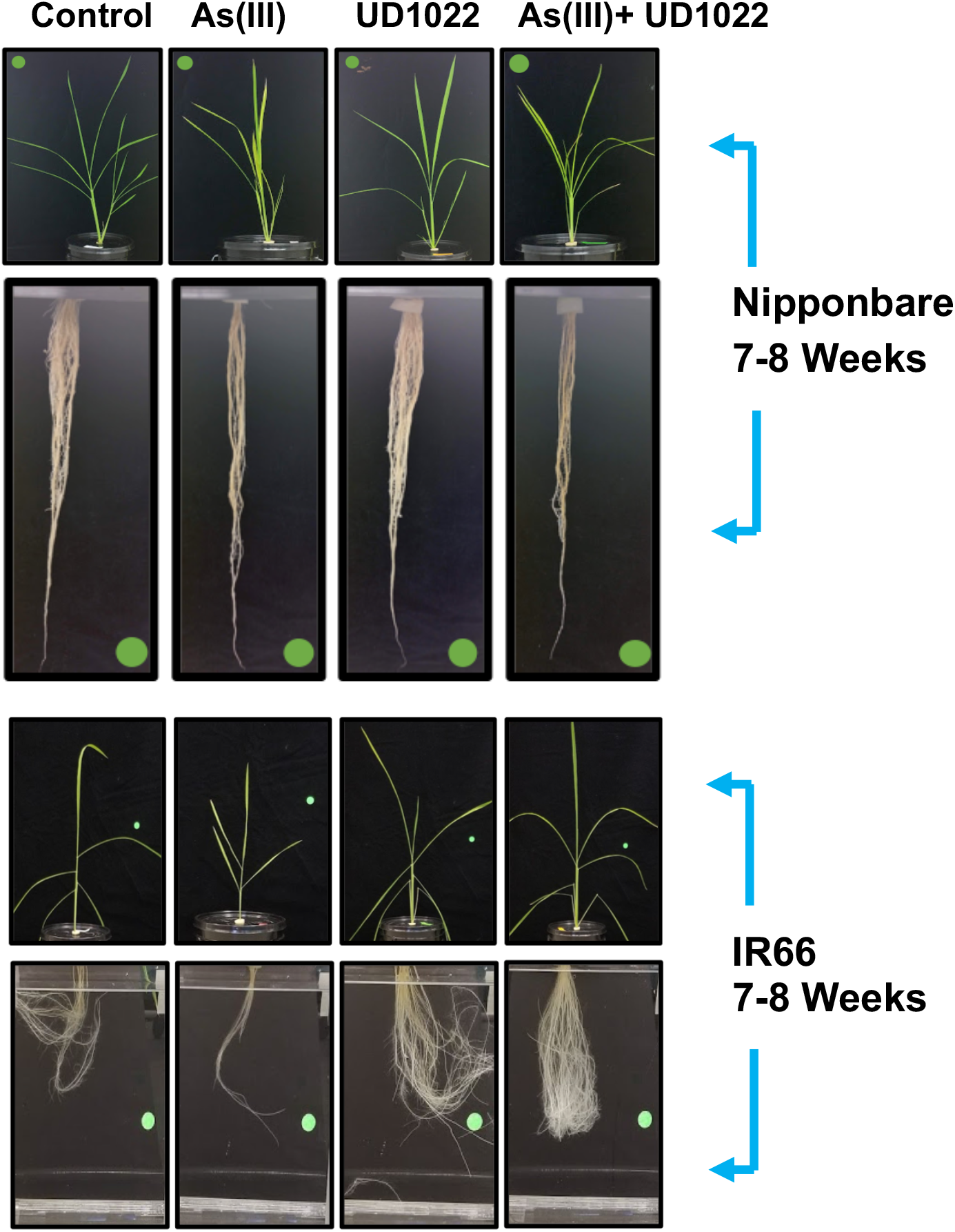
Nipponbare and IR66 rice plants inoculated with *B. subtilis* UD1022 and treated with a high concentration of As (~50 μM). Images show phenotypic differences in the shoots and roots of each plant. Representatives are shown at 7-8 weeks post inoculation and/or arsenic treatment. The green dot scales to 0.75 inches in diameter.

**Figure 8:**
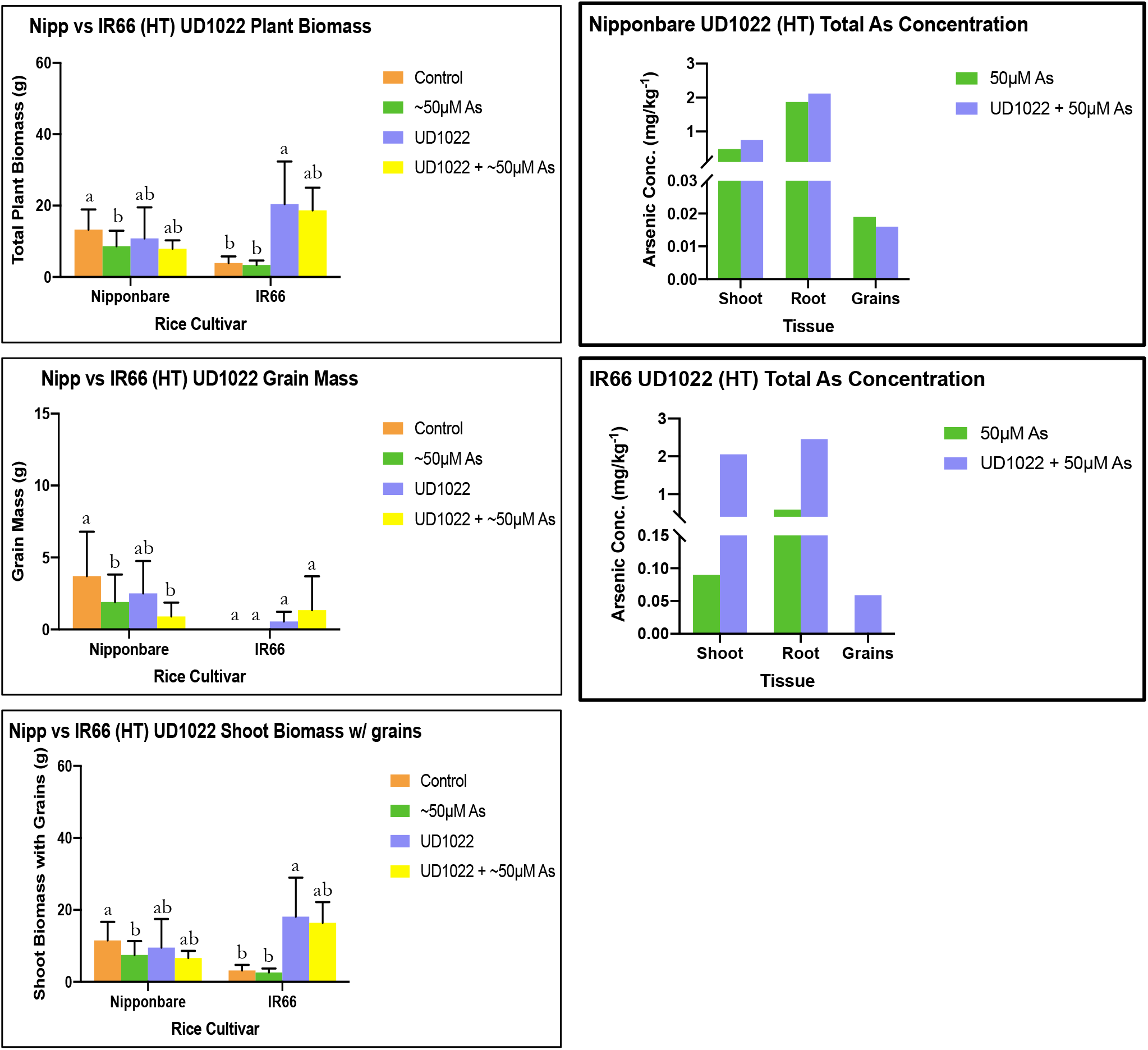
Total plant biomass for Nipponbare and IR66 rice exposed to As [III](~50 μM) and *B. subtilis* UD1022 bacteria. Means of common letters are not significantly different at *P* ≤ 0.05; according to Duncan’s Multiple Range Test.

Interestingly for As-susceptible IR66 plants treated with UD1022 and UD1022 + As showed increase in biomass over As treatments (Figures 7–8). As-uptake in UD1022 treatments showed that both Nipponbare and IR66 plants revealed similar As uptake (in shoots/roots tissues) patterns compared to other bacterial treatments (Figures 7–8). Interestingly, only UD1022 treated plants with As showed seed setting and grain formation in IR66 (Figures 7–8). These data suggest that a classical PGPR like UD1022 may provide minimal protection against high As treatment.

## Discussion

Arsenic has detrimental phytotoxic impacts on the aboveground physiology in rice, despite only having direct As contact with the roots. This supports previous data showing that abiotic stress in the form of As toxicity, no matter where it is applied, impacts the whole plant system (Lakshmanan et al., 2015). Photographs of the rice plants, taken throughout growth, visibly display the impacts of As on rice in high As environments. For both tolerant Nipponbare and susceptible IR66, the plant representative for the arsenic treatment displayed signs of As toxicity. These signs were visible earlier in the IR66 cultivar compared to Nipponbare. Rice plants grown in the hydroponics showed acute As toxicity both in shoot and root tissues. As-toxicity symptoms included stunted growth, chlorosis of leaves, disorganized plant structure, and reduced number of rice panicles. This is all consistent with previous studies showing the effects of As stress on rice yields (Moulick et al., 2018). The shoot and root phenotypes for Nipponbare and IR66 rice plants treated with As + EA104 or EA201 isolates presented signs of As recovery. The plants treated with EA104 and EA201 had relatively normal plant structure and the presence of grains compared to lone As treated plants that did not yield grain. It is rational to believe that EA201or EA104 may play a positive impact on mitigating arsenic uptake based on increased rice yields and past success of other *Arthrobacter sp.* being used for bioremediation of contaminated soils (Dsouza et al., 2015 and Krishnan et al., 2016).

When comparing the two rice cultivars, Nipponbare displayed more resistance against As stress than IR66 under EA106 and EA104 + As treatments. Nipponbare plants grown hydroponically with As showed less As accumulation in each tissue type. Surprisingly in the EA201 and UD1022 treatments, Nipponbare plants had an equal or increased amount of As in each tissue type compared to the IR66 plants with in the same trials. This could be a result of several factors; As uptake can be dictated by species availability and the plant growth stage (Awasthi et al., 2017). Since in our experiments As supplementation was done periodically to create a high As concentration, it is possible that As supplementation at a specific growth stage of plant may lead to more As uptake and mobility. We added As in the form of As(III), as it is known that As(III) is a more mobile and available As species for transport into the plant (Quaghebeur and Zdenko, 2003). It is not known how As uptake is modulated and facilitated under different growth stages in plants. It is speculated that As uptake in plants is facilitated when plants switch from vegetative to reproductive stage (Quaghebeur and Zdenko., 2003). Evaluating total As content and As speciation during multiple growth stages in plants under microbial inoculations may help understand what form of As is available and during which growth stages the plant is more vulnerable for increased As uptake.

The significant increase in total plant biomass and total grain yield in Nipponbare and IR66 rice plants exposed to EA104 and EA201 only could be attributed to the bacteria having better association with these rice varieties than the other tested isolates. Interestingly, both EA104 and EA201 were isolated from M-104 rhizospheric soil, and their association with Nipponbare and IR66 is not evaluated; at this juncture, we could only speculate that both EA104 and EA201 may have better colonization efficiency compared to EA104/EA201 under high As environment. The microscopy evidence with EA106 under high As treatment showed that increasing concentration of As may reduce EA106 colonization on rice roots. The ability of EA104/EA201 to tolerate high As concentration is also not known, and how it imparts its ability to colonize plants needs to be evaluated. The lack of microbial specificity to Nipponbare and IR66 could account for the variability between the tested plant varieties and the microbial isolate experiments. Our previous work showed that the treatment of EA106 and another natural rice isolate, *Pseudomonas chlororaphis* (EA105), against Nipponbare and a rice blast-susceptible Seraceltic rice variety exposed to low As environment resulted in low As accumulation in planta (Lakshmanan et al., 2015; 2016). In addition, we showed that EA106 treatment to Nipponbare plants supplemented with low As (5 μM) exposure, accumulated less As in shoots (Lakshmanan et al., 2015). Selecting natural rice rhizospheric microbes for attenuation of As uptake in our current and previous studies was based on the increased Fe-siderophore activity of the bacterial strains. It was argued that strains with increased Fe-siderophore activity may mobilize Fe on the root surface impeding As uptake by rice roots. Since our previous studies were performed under low As environment, we hypothesized that exposing rice plants with Fe-siderophore carrying strains may also protect plants against high As (50 μM) environment. It is clear from our study that microbe-mediated Fe-siderophore activity may not be the only mechanism which helps plants against As toxicity/uptake under high As toxicity environment.

The mechanisms that reduce As toxicity in plants are not very well understood. It is shown that various antioxidant molecules in plants attenuate As toxicity in plants, including rice (Brinke, et al., 2018). Rice plants exposed to As rich sediments showed overexpression of a few candidate genes such as UDPGT, pollenless3, and OsABI5 that may act as candidate gene clusters that could modulate plant resistance against As toxicity in rice (Brinke et al., 2018). Brinke et al., (2018) showed the exposure of rice plants to As-sediments also upregulated UDPGT (Os05t0527000–01) expression. The glucosyltransferase UDPGT (UDP-glucuronosyl/UDP-glucosyltransferase family protein) is linked to the molecular functions “quercetin 3-O-glucosyltransferase activity” and “quercetin 7-O-glucosyltransferase activity” (Brinke et al., 2018) and is involved in the anthocyanin pigment biosynthesis (Brinke et al., 2018). Anthocyanins belong to the group of flavonoids that are secondary metabolites known to be associated to As stress (Chakrabarty et al., 2009; Islam et al., 2015; Winkel-Shirley, 2002). The role of anthocyanins are also shown to attenuate other abiotic stresses such as aluminum toxicity (Winkel-Shirley, 2002). OsABI5, on the other hand, is known to be similar to ABA response element binding factor” or “*ABA-insensitive (ABI)5* gene”, which encodes a basic leucine zipper transcription factor known to regulate processes including pathogen defense, light and stress signaling as well as seed maturation and flower development in *A. thaliana* (Jakoby et al., 2002; Brinke et al., 2018). The role of other growth regulators such as auxin has also been reported to reduce As toxicity. Krishnamurthy and Rathinasabapathi (2013), showed that auxin transporters may help attenuate As toxicity in Arabidopsis. Krishnamurthy and Rathinasabapathi (2013), investigated the role of auxin transporters mutants such as *aux1*, *pin1* and *pin2*, which showed enhanced susceptibility against As(III) toxicity compared to the wild type (WT) plants. Auxin transport inhibitors significantly reduced plant tolerance to As(III) in the WT. Supplementation of exogenous indole-3-acetic acid (IAA) improved As(III) tolerance of *aux1* and not that of WT (Krishnamurthy and Rathinasabapathi 2013). How growth regulators such as auxin and ABA impact As transport and toxicity at the mechanistic level is not known.

Of late, the role of rhizosphere-mediated modulation for As uptake and biotransformation in soil has garnered more attention. The role of the microbiome and various microbial strains on As remediation and reduced As uptake in plants are now been investigated (Awasthi et al., 2017). Several α-Proteobacteria and β-Proteobacteria with enhanced arsenite (As(III)) oxidase gene (*aroA*-like) activity led to more As release and uptake in the plants and the rhizosphere (Jia et al., 2014). These results show that microbial processes in the rhizosphere may play an active role in As release and uptake by plants. The role of microbes that mediate and facilitate soluble Fe is also reported to modulate As release and uptake in the rice rhizosphere (Hashimoto and Kanke, 2018). How microbes and plants interact in an active manner with each other to either mobilize or abate As uptake needs to be elucidated.

In the future it would be interesting to isolate and test bacteria naturally occurring in the rhizosphere of Nipponbare and IR66 cultivars to evaluate if they have greater impacts on plant growth and abiotic resistance. Following the notion that As uptake can be greater at different times during rice development, perhaps adding additional doses of PGPRs near these developmental stages may mitigate As uptake into the plants (Awasthi et al., 2017). The fact that we administered microbes only once while adding more As in the hydroponics, also puts microbes at a disadvantage in our hydroponic system. Pulsed supplementation of microbes at a specific transition during a developmental stage of plants may also hold the key to attenuate As uptake by plants. To end, microbial intervention improves growth and yield in Nipponbare and IR66 rice varieties. We could not tease apart the relationship between establishment of microbes on the root surface to the increasing concentration of As in the hydroponic set up. Argumentatively, it may also be interesting to try multiple rice rhizospheric microbes under a consortium with low and high As envelopments with rice. The lack of synergism between microbes and the plant may also be because of the very high concentration of As in our studies impeding the microbial traits for As attenuation.

## Conclusion

The tritrophic (Rice-microbe -As) model system developed in our study showed that natural rice rhizospheric microbes provide minimal protection to the host plant under high As environment. The work presented also raises questions about the specificity of microbial inoculums to the plant hosts. Using a microbe (EA106) that showed protection against As under low As environment, we further showed that increasing As concentration may take the protection away from the host plant mediated by EA106. Though there is no direct evidence to increasing As concentration and microbial colonization on the root surface, we speculate that toxic levels of As may be microbicidal to disrupt efficient colonization of microbial inoculums for abating As uptake and toxicity.

## Data Availability Statement

The raw data supporting the conclusions of this article will be made available by the authors, without undue reservation.

## Conflict of Interest

The authors declare that the research was conducted in the absence of any commercial or financial relationships that could be construed as a potential conflict of interest.

## Author Contributions

All authors designed the experiments, edited and contributed to the final manuscript, and approved for submission. VGM completed lab work pertaining to hydroponic experiments and microbial treatments. VGM and HPB conceptualized the idea and designed the experiments.

## Funding

The work was supported by funding and support from University of Delaware and USDA NIFA to HPB (2016-67013-24846). The results, in part, are based on work reported in the master’s thesis of Victoria Gundlah-Mooney, which is freely available through the University of Delaware Library Institutional Repository (Gundlah-Mooney, V. (2019). Intervention of the rice microbiome in abating arsenic toxicity in rice (Order No. 27663789). Available from Dissertations & Theses @ University of Delaware. (2378066244).

## Notes

### Competing Interest Statement

The authors have declared no competing interest.

