## Supplementary material for "Rice rhizospheric microbes confer limited Arsenic protection under high Arsenic conditions": Figure S1

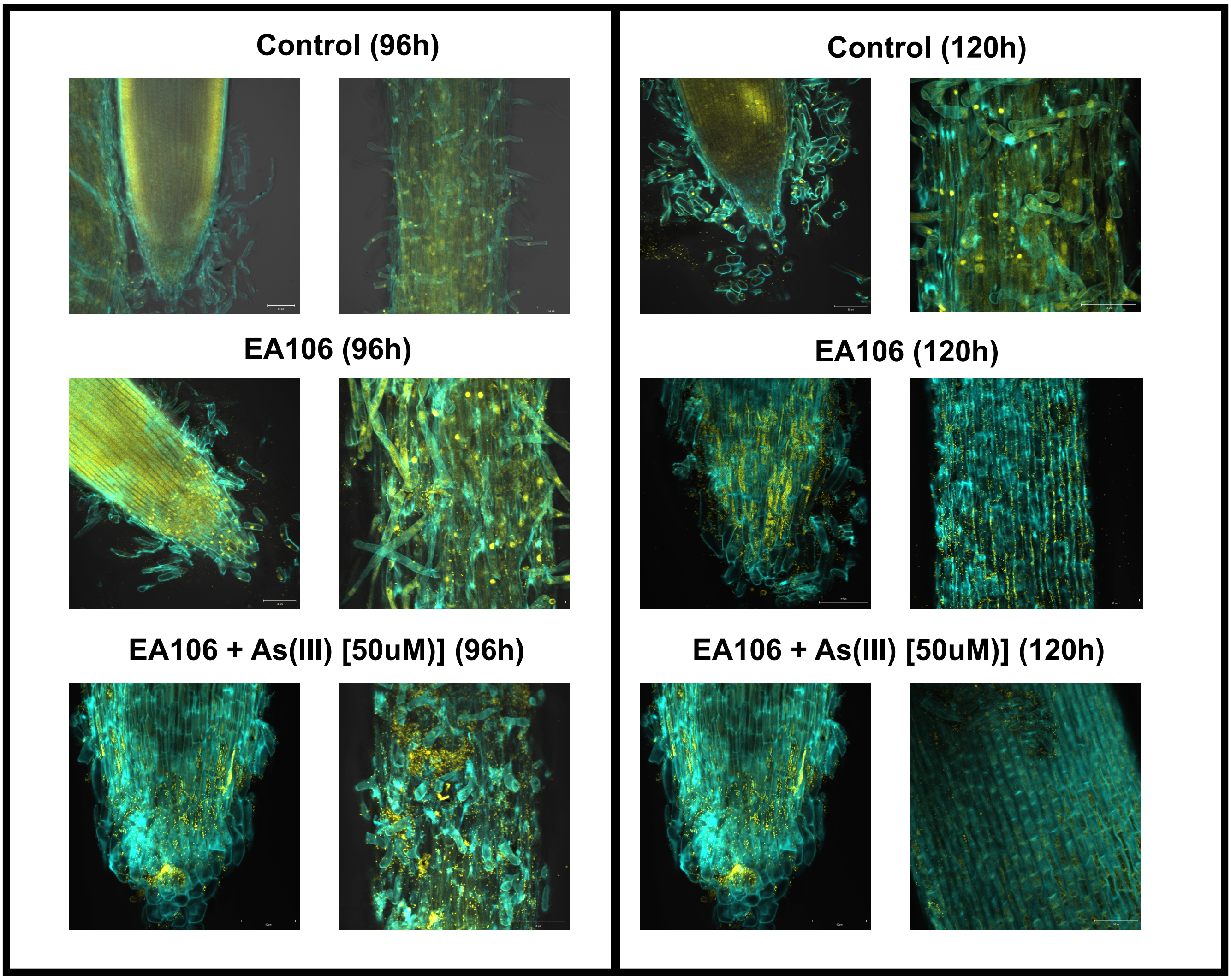


**Figure S1:** Micrographs depicting root colonization by *Pantoea* sp (EA106) in As-susceptible rice variety IR66. Micrograph of root tip and central elongation zone (*CEZ*) were taken post 96 and 120hr of microbial inoculum and As treatment. The yellow punctate fluorescence indicate bacterial presence and colonization on roots.
